# Rod signaling in primate retina: range, routing and kinetics

**DOI:** 10.1101/352419

**Authors:** William N Grimes, Jacob Baudin, Anthony Azevedo, Fred Rieke

## Abstract

Stimulus or context dependent routing of neural signals through parallel pathways can permit flexible processing of diverse inputs. For example, work in mouse shows that rod photoreceptor signals are routed through several retinal pathways, each specialized for different light levels. This light level-dependent routing of rod signals has been invoked to explain several human perceptual results, but it has not been tested in primate retina. Here we show, surprisingly, that rod signals traverse the primate retina almost exclusively through a single pathway, regardless of light level. Indeed, identical experiments in mouse and primate reveal large differences in how rod signals traverse the retina. These results require reevaluating human perceptual results in terms of flexible computation within this single pathway. This includes a prominent speeding of rod signals with light level – which we show is inherited directly from the rods photoreceptors themselves rather than from different pathways with different kinetics.

## Introduction

Rod photoreceptors contribute to vision across a million-fold range of light intensities. At the low end of this range - e.g. in starlight - photons are few and far between, and retinal circuits face the considerable challenge of detecting and reliably transmitting signals resulting from the absorption of individual photons (reviewed by (Field, Sampath, & Rieke, 2005)). Other challenges emerge as light levels increase. For example, in moonlight the high gain associated with detecting single photons, if unabated, would quickly saturate retinal responses. At the same time, the increased fidelity of light inputs in moonlight creates opportunities for more elaborate computations than possible in starlight.

A prominent hypothesis about how retinal circuits meet these changing demands proposes that the dominant route that rod-derived signals take through the retina depends on mean light level (Sharpe & Stockman, 1999; Bloomfield & Dacheux, 2001; Deans, Volgyi, Goodenough, Bloomfield, & Paul, 2002) - with different circuits specialized to handle the challenges associated with different light levels (Tsukamoto, Morigiwa, Ueda, & Sterling, 2001). These circuits include (Figure 1; reviewed by (Bloomfield & Dacheux, 2001; Field et al., 2005; Field & Chichilnisky, 2007)): 1) the primary pathway, in which dedicated rod bipolar cells transmit rod signals, 2) the secondary pathway, in which gap junctions convey rod signals to cones and the associated cone circuitry, and 3) the tertiary pathway, in which Off cone bipolar cell dendrites receive direct rod input. Work in rodent retina supports the hypothesized light level-dependent routing of rod signals (Soucy, Wang, Nirenberg, Nathans, & Meister, 1998; Deans et al., 2002; Trexler, Li, & Massey, 2005). In particular, while the primary pathway dominates responses of mouse ganglion cells at low light levels, the secondary pathway contributes substantially at intermediate to high rod light levels (i.e. mesopic conditions) (Deans et al., 2002; Ke et al., 2014; Grimes, Schwartz, & Rieke, 2014).

**Figure 1:**
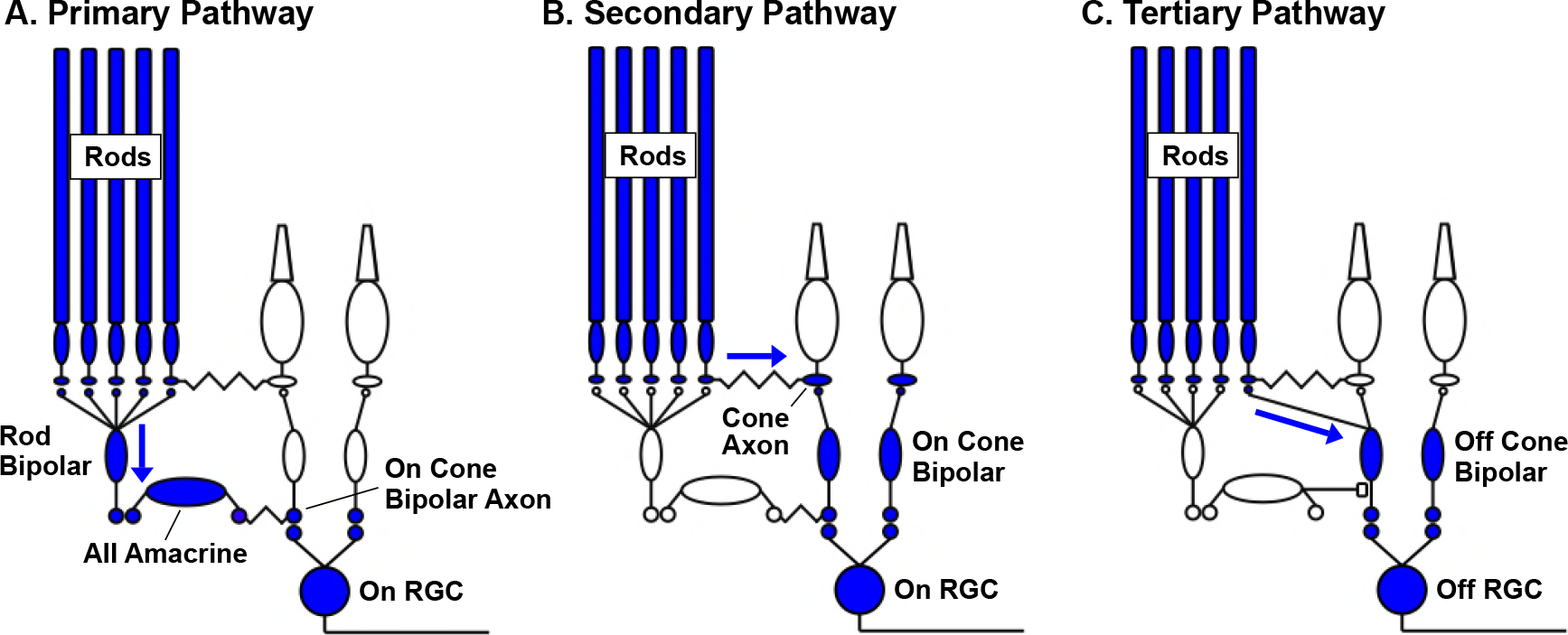
Rod signaling routing in the mammalian retina. A) In the primary pathway rod signals are routed through dedicated rod bipolar cells to AII amacrine cells. AII amacrine cells in turn transmit ‘On’ signals to On cone bipolar cells through dendro-axonal gap junctions and ‘Off’ signals to Off cone bipolar cells through glycinergic synapses (not shown, but see C). Cone bipolar signals are subsequently transmitted to retinal ganglion cells. B) In the secondary pathway, rod transmit signals via gap junctions to cone axons, and hence the associated cone circuitry (Kolb, 1977; Schneeweis & Schnapf, 1995b; Deans et al., 2002; Hornstein et al., 2005). C) In the tertiary pathway, rods transmit signals directly to Off cone bipolar cell dendrites (Soucy et al., 1998; Hack, Peichl, & Brandstatter, 1999; Tsukamoto et al., 2001b).

A light level-dependent routing of rod-derived signals has been invoked to explain several human perceptual observations (reviewed by (Sharpe & Stockman, 1999; Buck, 2004; Stockman & Sharpe, 2006; Buck, 2014; Zele & Cao, 2014)). For example, perceptual experiments show that the kinetics of rod-derived signals speed substantially as luminance levels increase from low to high mesopic conditions (Conner, 1982; Sharpe, Stockman, & MacLeod, 1989). Relatedly, differences in the kinetics of rod-and cone-derived signals play a central role in how these signals are combined perceptually (MacLeod, 1972; Frumkes, Sekuler, Barris, Reiss, & Chalupa, 1973; Grimes, Graves, Summers, & Rieke, 2015). The link between these perceptual phenomena and the routing of rod signals is based on the assumption that the primary pathway introduces larger delays in rod-derived signals than the secondary or tertiary pathways.

Despite its importance for understanding human vision, there are no direct tests of how rod-derived signals are routed through the primate retina. Here, we determine 1) the range over which rod photoreceptors control retinal output, 2) how and when rod signals traverse a given pathway, and 3) what mechanisms shape the kinetics of rod-derived retinal outputs. Surprisingly (and unlike mouse retina), we find that rod-derived signals in primate retina are largely restricted to the primary rod pathway even at mesopic light levels. Under the same conditions, the responses of rods themselves speed sufficiently to explain previous human perceptual results, indicating that this speeding does not require the rerouting of rod-derived signals through faster retinal pathways.

## Results

### Rod signaling range and saturation

We start by defining the range of light levels over which rod photoreceptors respond to light inputs and comparing this range to the range over which rod-derived signals are present in the retinal outputs.

We used suction electrode recordings to measure rod outer segment membrane currents. Figure 2A compares responses to a series of light steps (10 s) with the response to a saturating flash. Light steps producing ~40 R*/rod/s halved the outer segment current, and little dark current remained at 300 R*/rod/s. Figure 2B collects results from several such experiments. Mean backgrounds of 250 R*/rod/s suppressed 90% of the dark current (red line). This suppression was maintained for steps that persisted for 1-2 min (< 5% recovery of dark current, not shown).

**Figure 2:**
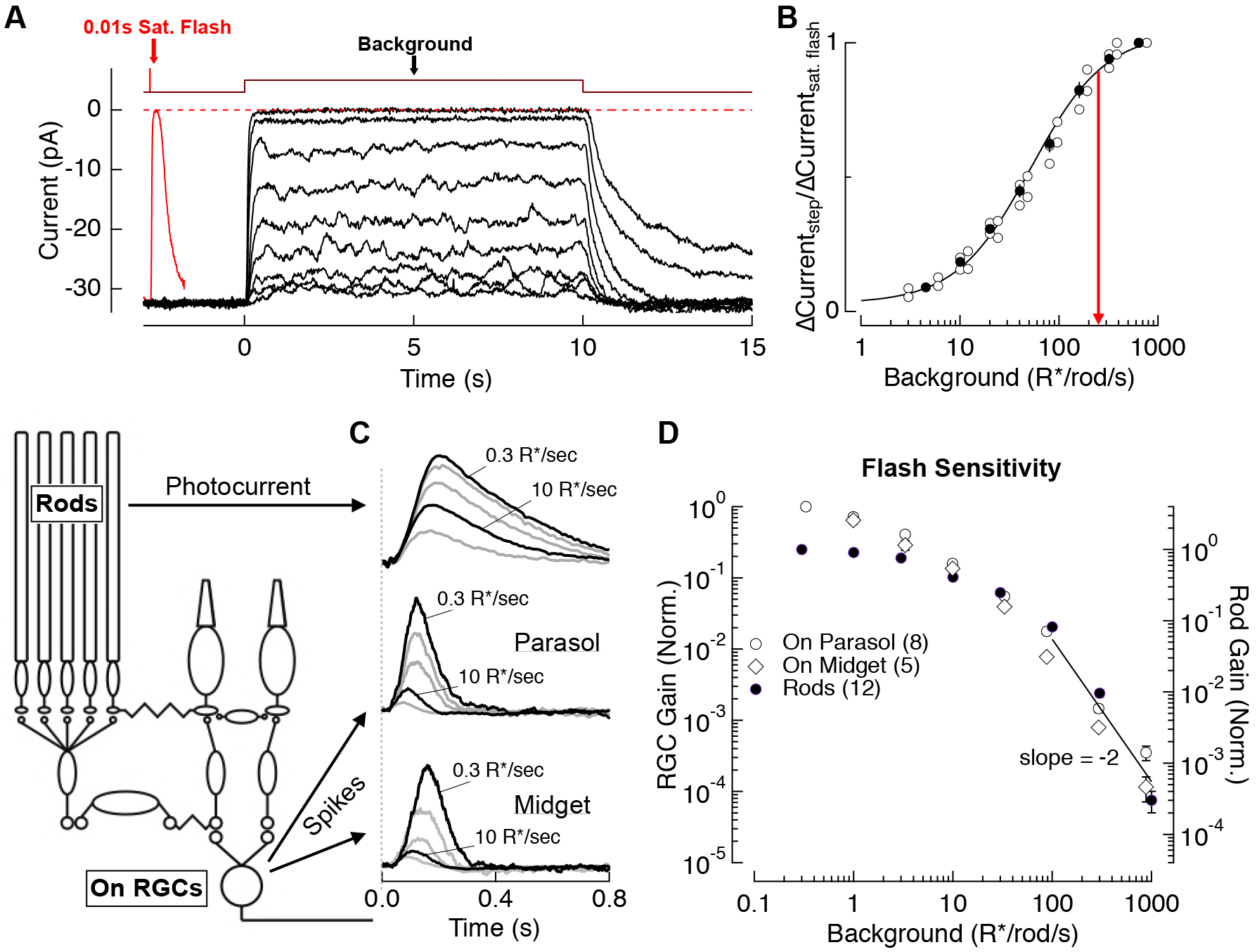
Evaluation of the range of rod signaling in phototransduction and retinal output. A) Direct recordings of photocurrent from rod outer segments (primate). Comparison of the responses to a short-wavelength saturating flash and a family of 10 s light steps or ‘backgrounds.’ B) Current suppression, relative to saturation, plotted as a function of background luminance. Red arrow (250 R*/rod/s) indicates 90% saturation. C) Gain of rod signals probed with short wavelength flashes on a range of rod adapting backgrounds in rod photoreceptors (top), On parasol RGCs (middle) and On Midget RGCs (bottom, traces are mean responses scaled by the flash strength in R*/rod, see Methods). D) Population data for rod gain measurements in rod photoreceptors and ganglion cells. Gain is normalized to that at 0.3 R*/rod/s. The solid line highlights the steep drop in gain associated with rod saturation.

To compare the range of rod signaling in rods with that in retinal ganglion cells (RGCs; the retinal output neurons), we measured rod photocurrents and RGC spike responses to brief short-wavelength flashes across a range of backgrounds (Figure 2C). Spectral measurements indicated that RGC responses to short-wavelength flashes were dominated by rods up to 100-300 R*/rod/s (see Methods). Figure 2D summarizes measurements of the background dependence of rod and RGC response gain (peak response divided by flash strength). Between darkness and 3 R*/rod/s, RGC gain declined considerably more than rod gain; this reduction in RGC gain reflects post-rod circuit adaptation (Lee, Pokorny, Smith, Martin, & Valberg, 1990; Purpura, Tranchina, Kaplan, & Shapley, 1990; Donner, Copenhagen, & Reuter, 1990; Dunn, Doan, Sampath, & Rieke, 2006; Dunn & Rieke, 2008; Schwartz & Rieke, 2013). For backgrounds between 3 and 100 R*/rod/s, both rod and ganglion cell gain declined approximately inversely with background, as expected for Weber’s Law (reviewed in (Rieke & Rudd, 2009)). Above this light level, rod signal gain in both rods and RGCs declined with increasing background more sharply than expected from Weber’s Law, falling by a factor of ~100 between 100 and 1000 R*/rod/s (black line). This abrupt decline in the gain of rod-derived signals coincides closely with the suppression of outer segment current (Figure 2B). The reduction in rod and RGC signal gain, like the reduction in rod outer segment current, was maintained for light steps that persisted for several minutes.

These experiments indicate that the primary limitation on the range of rod-derived signaling in the retinal output is a decrease in gain of rod signals, rather than decreased gain in post-rod circuitry. Rods may continue to weakly modulate retinal output signals above 300 R*/rod/s, but such responses are small compared to cone-derived responses for all but short wavelength stimuli (see Methods). Hence, we refer to the sharp drop in sensitivity above 300 R*/rod/s as rod saturation, and focus on rod signaling for backgrounds of 0 to 300 R*/rod/s.

### Rod signal routing

What route do rod signals take through the primate retina across the range of light levels identified above? The results described below show, surprisingly, that the rod bipolar pathway is the dominate route for rod-derived RGC responses across the full range of rod signaling.

We start by comparing responses of key cells in the primary and secondary rod pathways. One such cell is the AII amacrine cell. AIIs receive direct glutamatergic input from rod bipolar cells through AMPA-type glutamate receptors and make bidirectional electrical synapses with On cone bipolar cell axons (Figure 3A). Thus, under control conditions, AII responses reflect signaling from the primary pathway via rod bipolar cells and from the secondary pathway via gap junctional input from cone bipolar cells. In the presence of AMPA receptor antagonists, remaining AII responses reflect input only from the secondary pathway (Cohen, 1998; Trexler, Li, Mills, & Massey, 2001; Murphy & Rieke, 2008; Münch et al., 2009; Grimes, Schwartz, & Rieke, 2014; Ke et al., 2014). Because AMPA receptor antagonists likely alter retinal signaling in multiple ways, we performed complimentary experiments on horizontal cells that did not require pharmacological manipulation. We start by describing the AII results.

**Figure 3:**
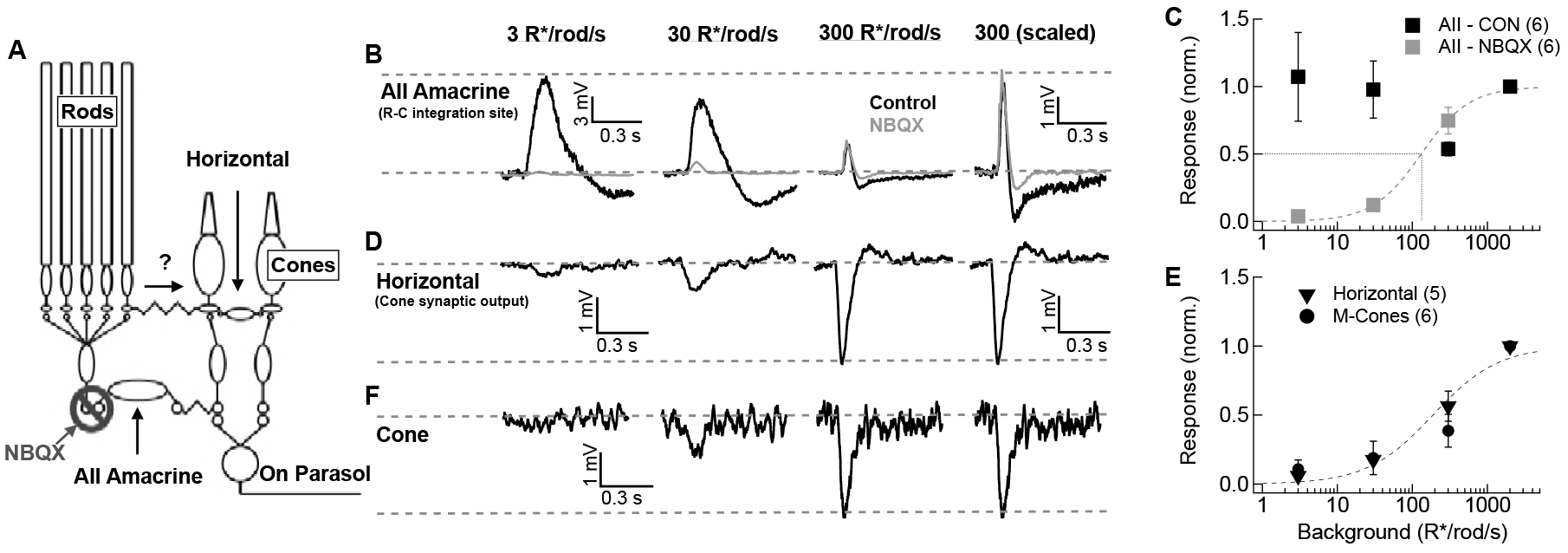
Comparison of rod signal strength in cells of the primary and secondary rod pathways. A) Schematic of the primary and secondary rod circuits and action of pharmacological manipulation used in the AII amacrine recordings. B,D,F) Responses to short wavelength 600% contrast flashes across a range of rod backgrounds in AII amacrine cells, H1 horizontal cells, and M-cones. B) Voltage responses of current-clamped AII amacrine cell before (black) and during exposure to NBQX (10 µM).Population data for AII amacrine recordings with and without NBQX. D) Voltage responses of current-clamped H1 horizontal cell. E) Population data for H1 horizontal and M-cone recordings. F) Voltage responses of an M-cone recorded in the perforated patch configuration.

After achieving a stable current-clamp recording, fixed-contrast short-wavelength flashes were delivered on backgrounds ranging from darkness to 2000 R*/rod/s (Figure 3B); these backgrounds extend beyond rod saturation (see Figure 2). The sensitivity of AII flash responses recorded in control conditions closely matched that of excitatory synaptic inputs to On parasol RGCs (Figure 3-figure supplement 1). This is consistent with the known role of AII amacrine cells in the circuits that convey rod signals to RGCs across light levels, and confirms that they are a reliable indicator of the strength of rod signals in On retinal circuits.

Block of AMPA receptors with NBQX dramatically altered AII responses – reducing responses more than 80% at backgrounds at or below 30 R*/rod/s (Figure 3B-C). Several properties of the NBQX-insensitive responses suggest that they originated in cones. First, they had shorter durations than control AII responses (Figure 3-figure supplement 1). Second, they grew with increasing background, reaching half-maximal amplitude at ~300 R*/rod/s. Most importantly for our purposes, NBQX-insensitive responses were largely absent over the range of light levels in which rods modulate the retinal outputs (i.e. below 300 R*/rod/s). This correspondence between rod saturation (Figure 2) and the emergence of NBQX-insensitive responses (Figure 3) suggests that the secondary rod pathway does not play a major role in transmitting rod signals to ganglion cells in primates, regardless of light level. This conclusion is, however, subject to the caveat that NBQX will impact retinal signaling in multiple ways.

To further test the hypothesized dominance of the primary rod pathway, we made current-clamp recordings from H1 horizontal cells. H1 horizontal cells receive direct synaptic input from L-and M- cones but not from rods (Figure 3A; (Kolb, 1970; Rodieck, 1998; Verweij, Dacey, Peterson, & Buck, 1999)); thus, these cells provide a readout of rod signals in the synaptic output of cones without the need for pharmacology. H1 responses, like the NBQX-insensitive responses of AII amacrine cells, were weak at low backgrounds but became pronounced at backgrounds ≥300 R*/rod/s (Figure 3D,E). In a subset of horizontal cell recordings, we also compared responses to short-wavelength (rod-preferring) and long-wavelength (cone-preferring) flashes (Figure 3 –figure supplement 2). At backgrounds below 300 R*/rod/s, responses to short wavelength flashes had substantially slower kinetics than responses to long wavelength flashes, consistent with previous work showing that rod-cone gap junctions can transmit rod signals to cones (Nelson, 1977; Schneeweis & Schnapf, 1995; Hornstein, Verweij, Li, & Schnapf, 2005). These responses, however, were small even for high contrast flashes. At backgrounds ≥300 R*/rod/s, responses to long-and short-wavelength flashes had very similar kinetics, suggesting that at these light levels both responses originated in the cones. Responses of cones were very similar to those of horizontal cells, with the emergence of a sizeable response only for backgrounds ≥300 R*/rod/s (Figure 3E,F). Rod responses measured in cones and horizontal cells in our experiments were similar in magnitude and kinetics to those in previous studies (Hornstein et al., 2005; Verweij et al., 1999) (Figure 3 – figure supplement 3).

The weak rod responses in cones and horizontal cells are consistent with the NBQX-insensitive responses in AII amacrine cells. The consistency of these results mitigates concerns about off-target effects of AMPA receptor block. Thus, these results collectively indicate that the secondary pathway plays, at most, a minor role in transmitting rod signals to ganglion cells in primate retina.

### Comparison of rod signal routing in mouse and primate

The dominance of the primary rod pathway in primate was unexpected, as previous experiments in rodents have assigned a substantial role to the secondary pathway in transmitting rod signals. To confirm this interspecies difference, we repeated several experiments from Figures 2 and 3 in mouse retina (Figure 4). Suction recordings from rod outer segments were again used to estimate the range of rod signaling. Responses of mouse rods saturated at higher light levels than primate rods (90% saturation at 1000 vs 300 R*/rod/s; Figure 4A). Voltage responses of AII amacrine cells in NBQX (Figure 4B) and horizontal cells in control conditions (Figure 4C-D) indicated that responses in the secondary pathway were half maximal at ~5 R*/rod/s, nearly two orders of magnitude lower than the half-maximal intensity of secondary-pathway rod signals in primate. Thus, the secondary pathway in mouse begins contributing to retinal output at light levels that are more than 100-fold below rod saturation, whereas in primate signals from the secondary pathway are largely absent below rod saturation.

**Figure 4:**
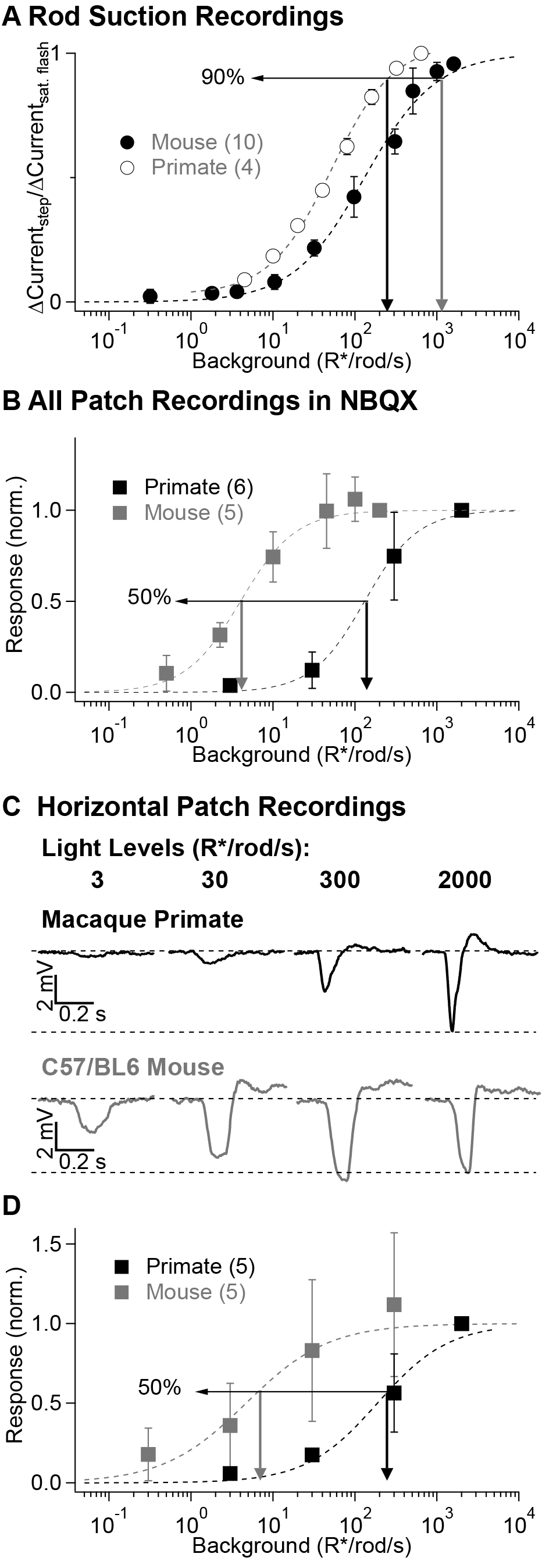
Comparison of secondary pathway activity in mouse and primate under identical experimental conditions. Steady-state measurements of the dependence of rod current on mean light level from suction recordings in primate and mouse as in Figure 2A, B) NBQX-insensitive responses in All amacrine cells recorded in current-clamp as in Figure 3, C) Horizontal cell current-clamp recordings to short wavelength flashes across a range of rod adapting backgrounds in primate (top) and mouse (bottom) as in Figure 3. D) Population data from horizontal cell recordings in primate and mouse.

### Routing of responses to continuous stimuli

We next considered whether continuous stimuli might more effectively elicit signals through the secondary pathway than the brief flashes used thus far. Sinusoidal stimuli are of particular relevance since they are used in many of the human perceptual studies that motivate the light level-dependent rod routing hypothesis. We started by recording excitatory synaptic input to an On parasol RGC in response to short-(rod-preferring) and long-(cone-preferring) wavelength sinusoidal stimuli at a mid-to-high mesopic light level (10 R*/rod/s). The contrasts of the short and long wavelength stimuli were adjusted so that when modulated individually at 2 Hz they produced roughly equal amplitude modulation of the RGC”s excitatory inputs. These contrast amplitudes were then held fixed as we explored responses to a range of temporal frequencies (Figure 5A,B). As observed previously (Dacey, Peterson, Robinson, & Gamlin, 2003), cone-derived responses were much larger than rod-derived responses at temporal frequencies ≥10 Hz (Figure 5B). We next recorded from H1 horizontal cells in the same piece of retina and measured responses to the same stimuli (amplitude and background) used in the parasol recordings. Across all temporal frequencies probed, the ratio of H1 responses to rod-and cone-preferring stimuli was at least 10 times smaller than that for On parasol responses to the same stimuli (Figure 5C,D). Hence, rod-derived signals in the secondary pathway are too weak to explain rod-derived RGC responses. Instead, the primary rod pathway dominates responses to both sinusoidal and flashed stimuli.

**Figure 5:**
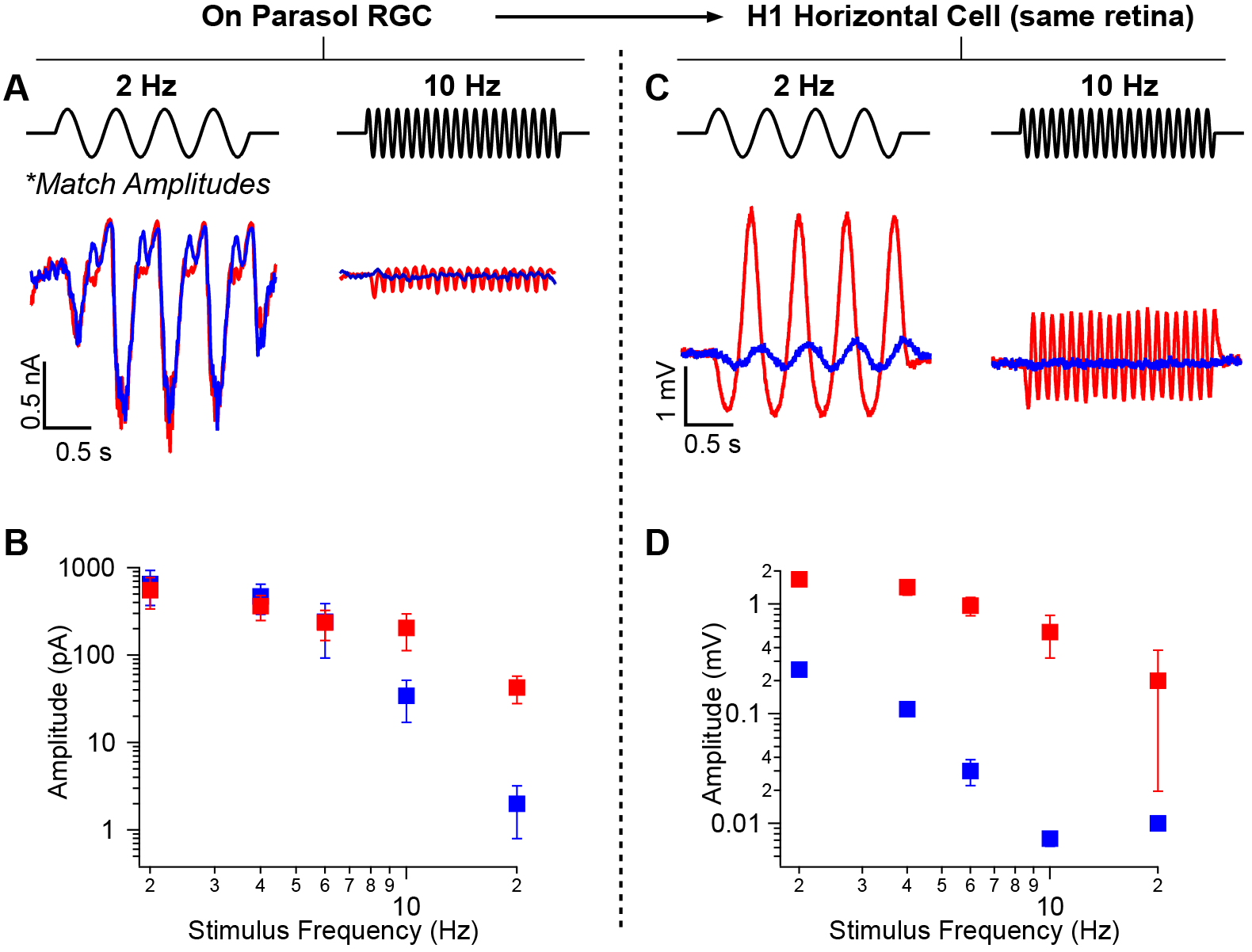
Rod-signals generated by sine-wave stimuli are also restricted to the RB pathway. A) Excitatory synaptic input recorded from an On parasol RGC in response to sinusoidally-modulated short (rod-preferring; blue traces) and long (cone-preferring; red traces) wavelength stimuli. Contrasts were adjusted to produce equal modulation at 2 Hz, and were then held fixed for all subsequent recordings (e.g. different frequencies, horizontal recordings). B) Population data from On parasol RGC recordings; response modulation versus stimulus frequency. C) Physiological response of an H1 horizontal cell (from the same retinal mount) to short and long wavelength stimuli for the same contrast used for the parasol cell in A. D) Population data from H1 horizontal recordings; response modulation versus stimulus frequency.

### Routing of rod signals in Off retinal circuits

Our results thus far suggest that the primary rod pathway continues to convey rod signals to On parasol ganglion cells even when the rods are approaching saturation. Does the primary pathway also dominate responses in Off retinal circuits? Rod signals could reach Off cone bipolar cells from three known sources (Figure 1): 1) dendritic input directly from rods (i.e. tertiary pathway), 2) dendritic input from cones (i.e. secondary pathway), and 3) axonal inhibitory input from the AII amacrine cell (i.e. primary pathway). The differences between On and Off circuits suggests that both the relative weighting of rod-and cone-derived signals and the routing of rod signals could differ.

We first compared the relative weighting of rod-and cone-derived signals in the responses of On and Off parasol and midget RGCs at a mean light level of 20 R*/rod/s (Figure 6A,B,C). Like the experiments in Figure 5, we began by adjusting the contrasts of rod-and cone-preferring stimuli so that they produced equal amplitude responses in an On parasol RGC (Figure 6A,C). After achieving a match, we presented these response-equated stimuli while recording from other RGC types in the same piece of retina. In Off RGCs, responses to rod-preferring stimuli were roughly half as large as responses to cone-preferring stimuli (Figure 6B,C). This indicates in turn that rod signals are less strongly routed through Off cone bipolar cells than On cone bipolar cells at this light level. If the secondary pathway (rod-cone electrical coupling) dominated, rod-and cone-derived signals would be mixed prior to transmission to On and Off bipolar cells, leading to similar weighting in On and Off circuits. Thus, this On/Off asymmetry supports our conclusion that the secondary rod pathway does not convey strong rod-derived signals in primate.

**Figure 6:**
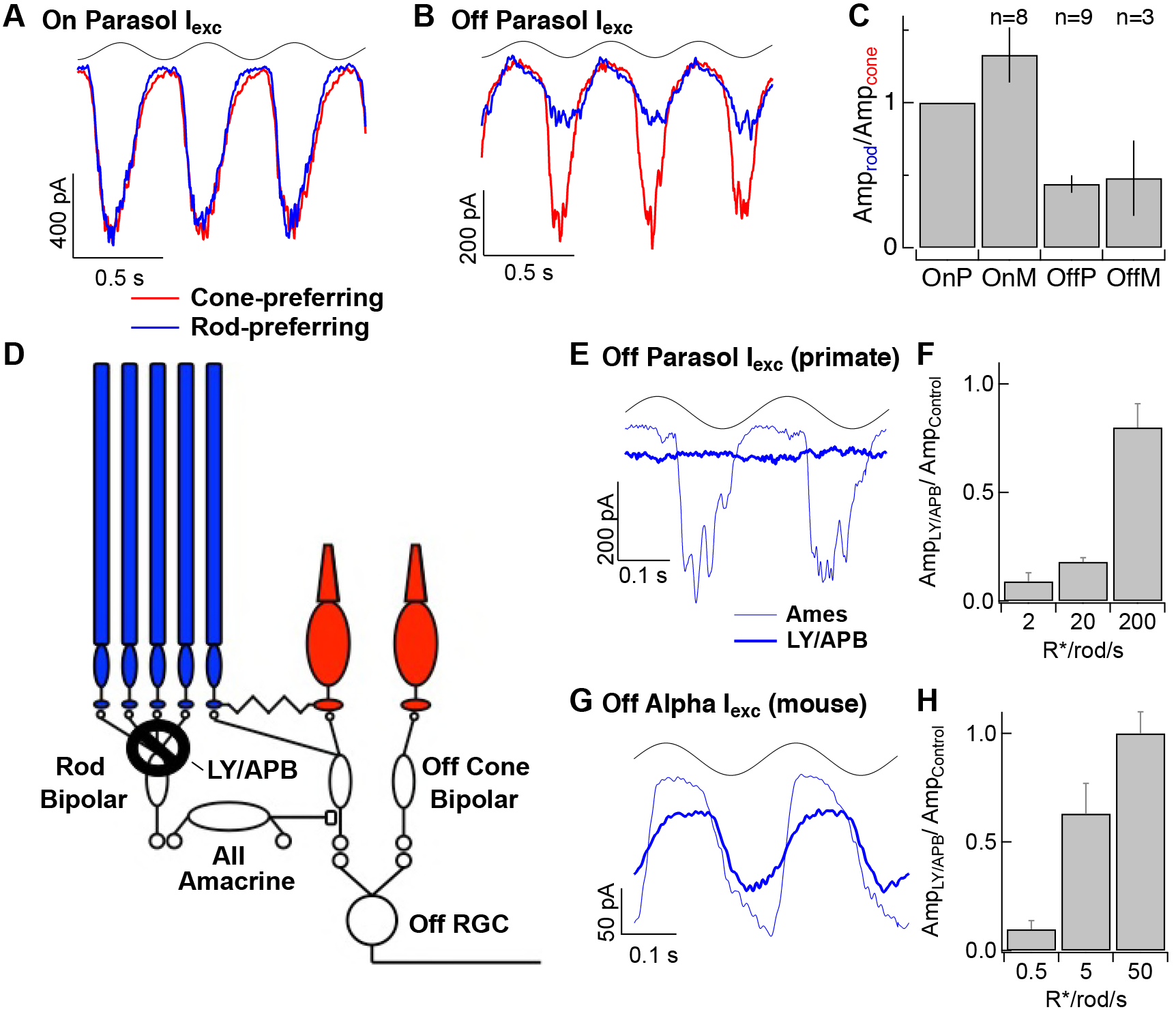
Rod signals in the tertiary rod pathway are weak. A) Excitatory synaptic input recorded from an On parasol RGC in response to sinusoidally-modulated short (rod-preferring; blue traces) and long (cone preferring; red traces) wavelength stimuli. Contrasts were adjusted to produce equal modulation at 2 Hz, and were then held fixed for subsequent recordings from other cell types (e.g. Off parasol RGC). B) Excitatory synaptic input recorded from an Off parasol RGC from the same retinal mount as A and in response to the same stimuli. C) Relative weighting of rod and cone signals in excitatory inputs to On and Off RGCs. D) Schematic of the primary and tertiary rod circuits that influence Off cone bipolar signaling and the actions of the mGluR6 agonist/antagonist mixture LY/APB. E) Rod signals (20 R*/rod/s) in excitatory inputs to an Off parasol in control conditions and after suppressing activity in all On bipolar cells with an mGluR6 agonist/antagonist cocktail (LY/APB, see Methods). F) Response ratio (cocktail:control) across cells as a function of mean luminance. Data are plotted as mean±SEM. G) Rod signals (5 R*/rod/s) in excitatory inputs to an Off alpha RGC (mouse) in control conditions and after suppressing activity in all On bipolar cells with an mGluR6 agonist/antagonist cocktail (LY/APB, see Methods). H) Response ratio (cocktail:control) across cells as a function of mean luminance.

Alternatively, rod-derived responses in Off RGCs could arise from the primary pathway via rod bipolar and AII amacrine cells or through the tertiary pathway via direct rod input to Off cone bipolar cells (Figure 6D). To distinguish between these possibilities, we used a mixture of an mGluR6 agonist (APB) and antagonist (LY341495) to suppress synaptic input to all On (both rod and cone) bipolar cells. This manipulation leaves intact both secondary and tertiary pathway input to Off cone bipolar cells. The agonist/antagonist mixture maintains the resting membrane potential of On bipolar cells and avoids anomalous changes in signaling that occur with a mGluR6 agonist alone (Ala-Laurila, Greschner, Chichilnisky, & Rieke, 2011).

Suppression of On pathways had little effect on the excitatory responses to long wavelength stimuli (Figure 6 – supplement figure 1), indicating, as expected, that direct cone input to Off cone bipolar dendrites remained intact. But suppression of On pathways reduced responses to short wavelength stimuli in Off parasol cells by more than 80% at backgrounds ≤20 R*/rod/s (Figure 6E,F). LY/APB reduced Off parasol excitatory synaptic input considerably less near rod saturation (20% reduction at 200 R*/rod/s). Thus, at light levels at or below 20 R*/rod/s, most of the excitatory input to an Off parasol cell requires activity of On bipolar cells, while above 200 R*/rod/s this is not the case. The sensitivity of Off parasol responses to suppression of signaling on On bipolar cells indicates that rod signals reach Off RGCs mainly through the primary rod pathway throughout the mesopic range.

We repeated the LY/APB experiments in mouse retina to provide a direct comparison across species (Figure 6G,H). At a background of 0.5 R*/rod/s, LY/APB reduced excitatory inputs to Off alpha RGCs by ~90%, indicating that the primary pathway dominates signaling at this light level. At a background of 5 R*/rod/s, LY/APB reduced responses by 35%, and at a background of 50R*/rod/s, responses were little affected. This is consistent with the results from Figure 4 that show that sizable rod signals reach AII amacrine cells through On cone bipolar cells in mouse retina, and correspondingly that the secondary pathway conveys significant rod signals at mesopic light levels.

Collectively, the experiments in Figures 3-6 indicate a surprising difference between how rod signals traverse the mouse and primate retinas. Specifically, unlike the situation in mouse, the primary rod pathway provides the dominant route that rod-derived signals take through the primate retina across scotopic and mesopic light levels. This dominance of the primary pathway means that perceptual phenomena previously attributed to a change in routing need to be reinterpreted (e.g. section on kinetics below); it also maximizes opportunities to independently process rod and cone signals since they are not mixed until late in the retinal circuitry.

### Rod signal kinetics

Perceptual experiments show that the kinetics of rod-derived signals speed relative to cone-derived signals as light levels increase (Sharpe et al., 1989). This speeding is often attributed to a luminance-dependent change in the dominant route that rod-derived signals take through the retina (reviewed by (Buck, 2004; Sharpe & Stockman, 1999; Stockman & Sharpe, 2006; Buck, 2014)), but the experiments described above suggest that such rerouting does not occur. Instead, as described below, the shift in kinetics of rod-derived signals appears to originate within the rods themselves.

We first determined whether responses of RGCs under our experimental conditions exhibited kinetic shifts similar to those observed perceptually. Spikes (Figure 7A) and excitatory synaptic inputs (Figure 7B) were recorded from On parasol RGCs in response to sinusoidal modulation across a range of light levels (0.2-100 R*/rod/s) over which rods dominate RGC responses. Both spike responses and excitatory synaptic inputs sped by ~100 ms across the range of light levels probed (Figure 7C). This speeding did not depend strongly on temporal frequency (data not shown). Kinetic changes between 1 and 10 R*/rod/s were similar in magnitude to those observed perceptually (Sharpe et al., 1989), suggesting a common origin in retinal circuits. The conserved routing of rod signals across these light levels indicates that these kinetic shifts occur within elements of the primary rod pathway rather than from a change in the dominant pathway conveying rod signals through the retina.

**Figure 7:**
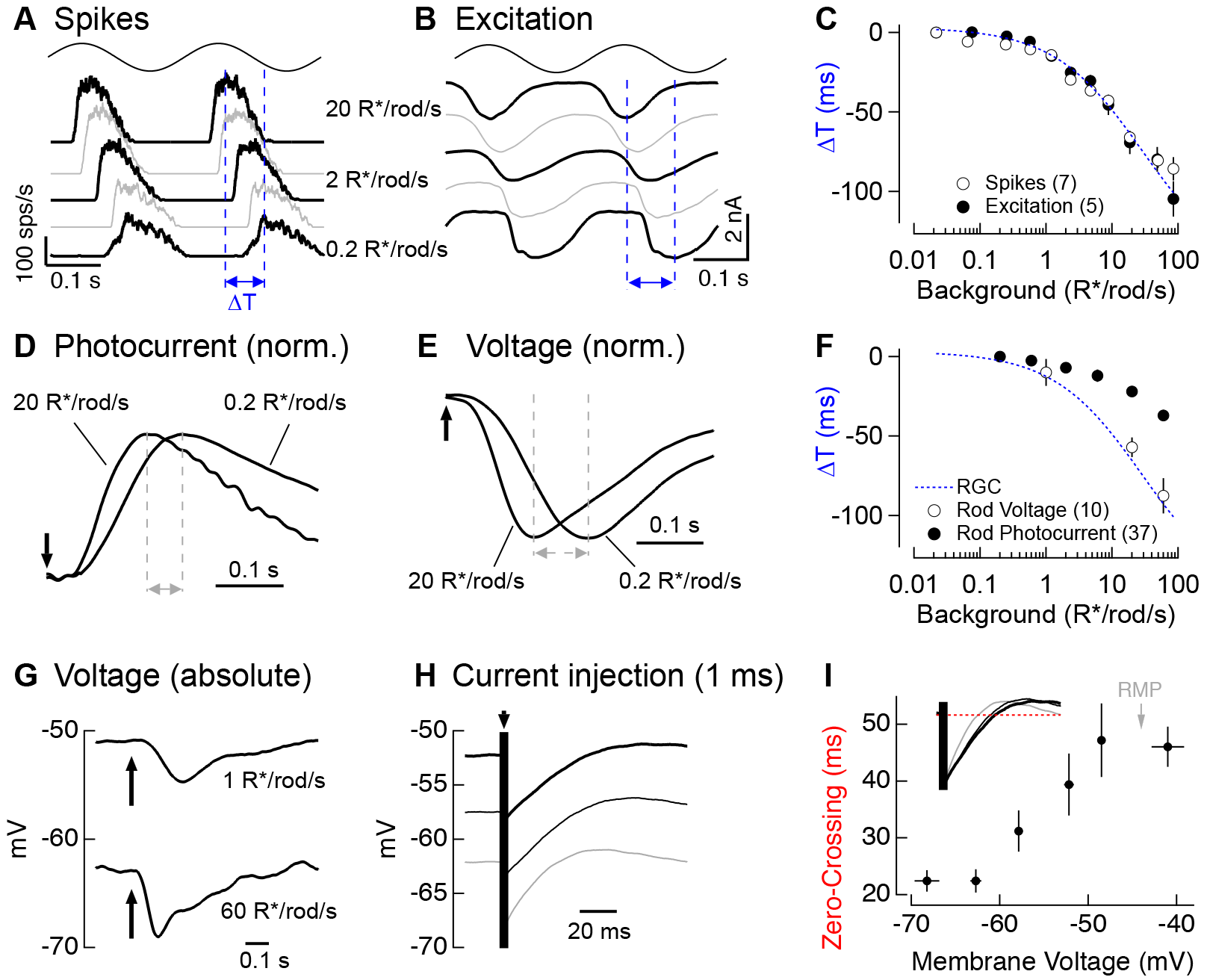
Change in kinetics of rod-derived responses across light levels. A) Spike responses to 4 Hz sinusoidal stimuli recorded from an On parasol across a range of rod backgrounds. B) Excitatory input currents to the same 4 Hz stimuli. C) Temporal shifts in On parasol responses plotted versus background luminance. D) Recordings of rod outer segment photocurrents in response to brief flashes across a range of rod backgrounds (normalized). E) Whole-cell current-clamp recordings from rod photoreceptors to brief flashes across backgrounds (normalized). F) Temporal shifts in rod photoreceptor recordings plotted versus rod background. G) Exemplary current-clamp recording of a rod photoreceptor responding to a brief flash on two different rod backgrounds. Rods exhibited large steady-state hyperpolarizations as luminance levels increased. H) Membrane response of a rod to a 1 ms current injection recorded at various physiological membrane potentials. I) Measured zero-crossing time of rod responses to 1 ms current injections plotted as a function of baseline membrane potential.

We returned to rod suction recordings to determine if phototransduction contributes to the kinetic shifts in the retinal output. Measuring rod responses to sinusoidal stimuli for light levels below 1 R*/rod/s would be near impossible due to the noise associated with stochastic photon absorption. Instead, we measured responses to brief flashes (Figure 7D), and used these to infer responses to sinusoids (see Methods). Flash and inferred sinusoidal responses sped as light levels increased (Figure 7F), but not to the full extent observed in responses of RGCs. Specifically, the change in kinetics observed in rod phototransduction currents could account for ~40% of the change in kinetics observed in RGC responses (Figure 7F, compare dashed blue line and closed circles).

Recent work on current-to-voltage transformations in mouse and goldfish photoreceptors indicates that intrinsic conductances can speed visual responses in rods and cones (Sothilingam et al., 2016; Seeliger et al., 2011; Della Santina et al., 2012; Howlett, Smith, & Kamermans, 2017). Could a similar mechanism account for the additional speeding between rod outer segment currents and RGC responses? To answer this question, we made whole-cell current-clamp recordings from rod photoreceptors in retinal slices. To avoid rundown of responses we focused on a limited number of backgrounds. Rod voltage responses (Figure 7E) to brief flashes showed larger changes in kinetics than current responses (Figure 7D). Further, when we used the flash responses to predict responses to sinusoidal stimuli, we found that they could fully explain the ~100 ms change in kinetics of RGC responses to sinusoidal stimuli (Figure 7F, dashed blue line and open circles).

The speeding of the rod voltage response coincided with a large steady-state hyperpolarization of the rod membrane potential (9.0± 0.7 mV upon a step from 1 to 60 R*/rod/s, mean ± SEM, n=10; Figure 7G). To test whether this hyperpolarization could alter the rod membrane time constant, we measured responses to brief hyperpolarizing current pulses across the physiological range of membrane voltages (with the voltage set by injecting steady currents; Figure 7H). Hyperpolarization indeed sped the membrane time constant by a factor of ~2 (Figure 7I), and this effect was well-matched to the physiological range of rod voltages (resting membrane potential in darkness of −44 ± 1 mV, mean ± SEM, n=10).

These experiments indicate that the shifts in kinetics of the rod-derived retinal outputs are largely inherited from the rods themselves rather than a light-dependent shift in the routing of rod signals through fast and slow retinal circuits. The speeding of rod responses can be explained by changes in the kinetics of phototransduction together with voltage-dependent changes in the conversion of outer segment currents to voltages.

## Discussion

### Meeting the challenges of mesopic vision

Sensory systems face the considerable challenge of encoding and processing widely varying inputs, and retinal processing across the 24-hour light-dark cycle exemplifies these challenges. Near visual threshold (e.g. starlight), photons arrive at individual rods only once every hundred to thousand seconds on average. Reliably detecting these sparse inputs requires integration of signals across many rods and across time. Under mesopic conditions (e.g. dawn or dusk), however, the rate at which photons arrive at the retina increases as much as one million-fold; this increased rate of photon arrivals alters the functional challenges and opportunities facing retinal circuits. For example, controlling the gain of rod-derived responses to avoid saturating retinal responses becomes a dominant requirement, and the improved signal-to-noise ratio of the inputs enables computations that are not possible in starlight.

A long-standing hypothesis about how rod vision operates effectively from starlight to twilight is that rod signals are routed through different retinal circuits under different conditions, such that the amplification and filtering properties of the dominant circuit are matched to the properties of the input (Tsukamoto, Morigiwa, Ueda, & Sterling, 2001). Thus, mechanisms in the primary or rod bipolar pathway are well suited to amplify single photon responses and remove noise (Field & Rieke, 2002; Grimes, Hoon, Briggman, Wong, & Rieke, 2014) and hence support vision in starlight. As light levels increase, the gain of rod signals measured in the retinal output decreases considerably (Lee et al., 1990; Purpura et al., 1990; Donner et al., 1990; Dunn et al., 2006; Dunn & Rieke, 2008; Schwartz & Rieke, 2013). RGC responses also speed considerably with increasing light level (Figure 8A). The result is a matching of temporal integration (Figure 8B) and signal-to-noise ratio (Figure 8C) to the statistical fluctuations in the light input.

**Figure 8:**
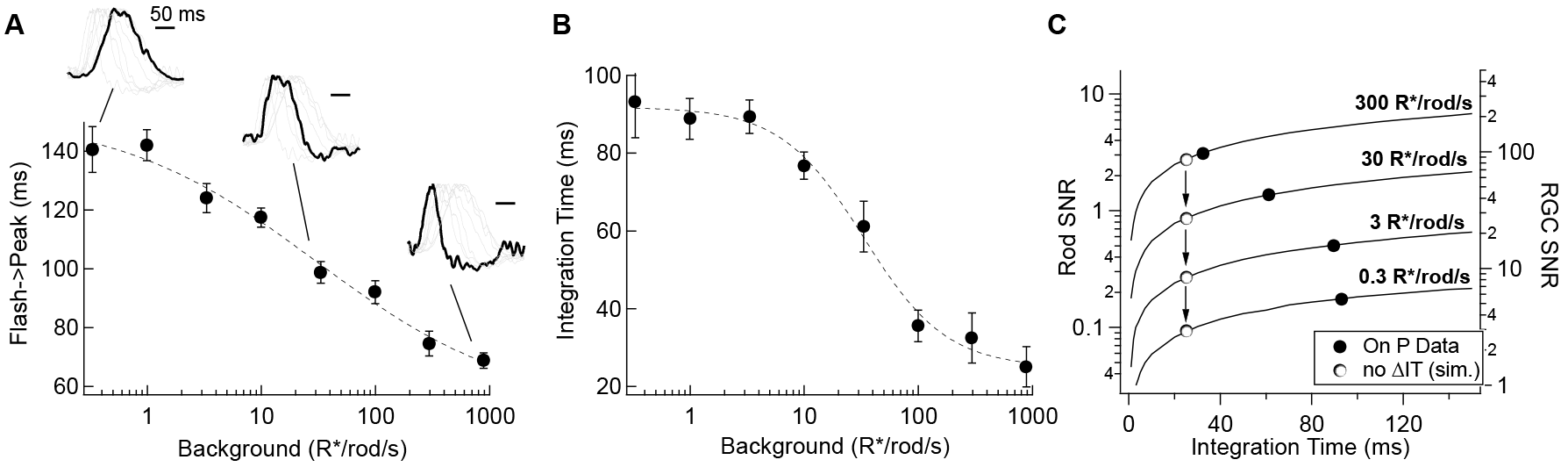
Changes in rod kinetics: tradeoffs between timing and SNR. A) The time-to-peak of rod-derived retinal outputs is reduced as rods adapt to brighter backgrounds. The time-to-peak of rod-derived signals in RGCs plotted as a function of background luminance. Inset traces show normalized PSTHs of spike responses from On parasol RGCs, with three luminance levels (i.e. 0.3, 30 and 900 R*/rod/s) highlighted. B) Integration time of retinal output measured from On parasol RGCs as a function of background luminance. C) Longer integration times improve the reliability of retinal encoding. Solid lines represent the SNRs experienced by individual rods (left axis) and individual RGCs (right axis) as a function of integration time for a range of photon capture rates (i.e. luminance). Black markers represent integration time measurements taken from On parasol RGC spike recordings. White markers represent a simulated scenario in which integration time does not increase as luminance decreases (i.e. integration time observed at the highest background tested is held constant).

This speeding of rod signals has been attributed to a shift in routing of rod signals from the (presumed slow) primary pathway to the (presumed fast) secondary or cone bipolar pathway. Experiments in non-primate retinas, particularly mice, support this hypothesis (Soucy et al., 1998; Deans et al., 2002; Trexler et al., 2005). The work described here shows, unexpectedly, that rod-derived signals in primate retinal ganglion cells are dominated by the primary pathway across light levels. This implies that light level-dependent changes in rod signaling in primate are not due to a change in routing, but instead to flexibility within the primary pathway. Specifically, we find that the speeding of rod signals in the retinal output can be explained by a change in kinetics of signals in the rods themselves - with approximately equal contributions from phototransduction and inner segment conductances. Inner segment conductances similarly speed responses of mouse rods and goldfish cones (Sothilingam et al., 2016; Seeliger et al., 2011; Della Santina et al., 2012; Howlett et al., 2017).

### Segregation of rod and cone signals

Recent anatomical and functional studies highlight differences in the segregation of rod and cone signals between rodent and primate retina. Specifically, mouse rod bipolar cells, long thought to receive input exclusively from rods, make some contacts with cones (Behrens, Schubert, Haverkamp, Euler, & Berens, 2016) and can convey cone-derived signals (Pang et al., 2010; Szikra et al., 2014). Cone inputs to rod bipolar cells have not been observed in primate retina. Similarly, rod contacts on cone bipolar dendrites appear more numerous in mouse than primate (Tsukamoto & Omi, 2014; Tsukamoto & Omi, 2016). This suggests greater mixing of rod and cone signals in rodent retina than in primate retina.

Our work here shows a surprisingly clear separation of rod and cone signals in primate retina – with even the well-established gap junctions between rods and cones (Kolb, 1977; Schneeweis & Schnapf, 1995; Hornstein et al., 2005) conveying at most a small rod-derived signal. The resulting separation of rod and cone signals through most of the retinal circuitry maximizes opportunities for independent processing. Such independent processing may be particularly important under mesopic conditions when rod signals are approaching saturation and cone signals are small and threatened by noise. Our work also provides a cautionary reminder that mechanistic studies derived from mouse may not be directly applicable to humans.

### Rod Saturation

Rod suction and RGC recordings (Figures 2 and 4) indicate that rods are largely saturated at ~300 R*/rod/s in primates and at ~1200 R*/rod/s in mice. Our physiological estimate of the upper end of rod signaling agrees well with human perceptual results (Sharpe, Fach, & Stockman, 1992). However, recordings from rodents reveal that rods can modulate retinal outputs at considerably higher light levels (Yin, Smith, Sterling, & Brainard, 2006; Naarendorp et al., 2010). These remaining rod responses rely at least in part on a form of bleaching adaptation that allows rods to regain sensitivity after prolonged exposure (>10 min) to photopic backgrounds (Tikidji-Hamburyan et al., 2017). On short time scales (≤10 min) and consistent with our results (Figure 4), Tikidji-Hamburyan and colleagues found that rods could no longer modulate RGC output at backgrounds at or above 104 R*/rod/s. It will be interesting to see if slow adaptation mechanisms found in mouse rods are also present in primates.

### Linking neural circuits to perception

Interactions between rod and cone signals affect many aspects of mesopic vision (reviewed by (Buck, 2014; Stockman & Sharpe, 2006)). An important factor controlling these interactions is the relative timing of rod and cone signals. This relative timing shifts considerably between low and high mesopic light levels as rod signals speed relative to cone signals (Sharpe et al., 1989). The experiments described here indicate that the speeding of rod-derived signals is not due to a change in routing, as often assumed, but instead occurs largely within the rods themselves. Thus, this work implicates rod adaptation, and not rerouting, as the critical retinal mechanism that shapes the time course of rod-derived signals and ultimately human perception under scotopic and mesopic conditions.

## Methods

### Electrophysiology

Experiments were conducted on whole mount or slice (200 μm thick) preparations of primate (rhesus macaque) or mouse (C57/BL6) retina as previously described (Dunn, Lankheet, & Rieke, 2007; Trong & Rieke, 2008). Retinas were obtained through the Tissue Distribution Program of the Washington National Primate Research Center; all procedures followed protocols approved by the Institutional Animal Care and Use Committee at the University of Washington. In brief, pieces of retina attached to the pigment epithelium were stored in ~32-34° C oxygenated (95% O_2_/5% CO_2_) Ames medium (Sigma) and dark-adapted for >1 hr. Pieces of retina were then isolated from the pigment epithelium under infrared illumination and either flattened onto polyL-lysine slides (whole mount: cone, horizontal cell, AII amacrine cell and RGC recordings) or embedded in agarose and sliced (rod recordings). Once under the microscope, tissue was perfused with Ames medium at a rate of ~8 mL/min.

Extracellular spike recordings from On parasol retinal ganglion cells used ~3 MΩ electrodes containing Ames medium. Voltage-clamp whole-cell recordings used electrodes (RGC: 2-3 MΩ, AII, HC: 5-6 MΩ) containing (in mM): 105 Cs methanesulfonate, 10 TEA-Cl, 20 HEPES, 10 EGTA, 2 QX-314, 5 Mg-ATP, 0.5 Tris-GTP and 0.1 Alexa (488) hydrazide (~280 mOsm; pH ~7.3 with CsOH). Current-clamp whole-cell recordings were conducted with electrodes (AII: 5-6 MΩ, HC: 5-6 MΩ, rods, cones: 10-12 MΩ) containing (in mM): 123 K-aspartate, 10 KCl, 10 HEPES, 1 MgCl_2_, 1 CaCl_2_, 2 EGTA (omitted for rod and cone recordings), 4 Mg-ATP, 0.5 Tris-GTP and 0.1 Alexa (488) hydrazide (~280 mOsm; pH ~7.2 with KOH). In initial whole-cell experiments, cell types were confirmed by fluorescence imaging following recording. To isolate excitatory synaptic input, cells were held at the estimated reversal potential for inhibitory input (~-60 mV). This voltage was adjusted for each cell to maximize isolation. Absolute voltage values have not been corrected for liquid junction potentials (K+-based = −10.8 mV; Cs+-based = −8.5 mV).

Perforated patch clamp recordings from cone photoreceptors were performed in current clamp using electrodes (9-11 MΩ) containing (in mM): 115 potassium aspartate, 1 MgCl_2_, 4 KCl, 10 HEPES, 10 diTris phosphocreatine hydrate, 4 Mg-ATP, 0.5 Tris-GTP (276-278 mOsm with potassium aspartate, pH 7.1-7.15 with KOH). Gramicidin was added to the internal solution at 30μg/mL. Upon sealing on a cell, access was monitored by tracking the membrane potential as well as the response amplitude to a constant amplitude probe flash. Once access equilibrated (~5 - 25 minutes), recordings were started. Throughout perforated patch recordings, the membrane potential and response to a reference flash were monitored to ensure that electrical access to the cell remained stable.

For suction recordings, a suspension of finely chopped retina was transferred to a recording chamber, pieces of retina were briefly allowed to settle, then perfused (2-3 ml/min, 32 ± 1 °C). Suction electrodes (3-4 MΩ, tip inner diameter of ~1.6μm) were filled HEPES-buffered Ames and were voltage clamped at 0 mV. Individual rod outer segments were drawn into the recording pipette under gentle suction.

Activity of mGluR6 receptors of On bipolar cells was suppressed in some experiments using a mixture of LY341495 (2.5 μM) and APB (7.5 μM). This approach was chosen to suppress modulated responses of On bipolar cells while minimizing perturbations to the state of the retina associated with tonic changes in mGluR6 receptor activity (Ala-Laurila et al., 2011).

### Cell selection criteria

Prior to recording data from all cells except photoreceptors, the sensitivity of a given piece of retina was estimated from responses of On parasol RGCs. Two criteria were used to determine whether to continue with data collection (Ala-Laurila and Rieke, 2014; Grimes, Graves, Summers, & Rieke, 2015): (1) high sensitivity when fully dark-adapted, as measured by a clear increase in spiking to flashes producing ~0.002 R*/rod; (2) clear cone-dominated responses to flashes from 640 nm LED at backgrounds of 20 R*/rod/s.

All reported data from horizontal cells, AII amacrine cells and RGCs was from retinas in which these criteria were met. In suction electrode recordings, rod photoreceptors were selected for recording and analysis if their outer segment had not been obviously damaged by the suction procedure and if they had maximal light responses exceeding 20 pA for primate and 8 pA for mouse. In whole-cell recordings, rods were retained for analysis if they exhibited > 8 mV changes in resting potential for light steps from 1 to 60 R*/rod/s. In all cases, data from any recorded cells in which these criteria were met was retained and reported in relevant figures.

We collected data generally from 5-10 cells for each experimental manipulation; this target population size was based on checking similarity of effects across cells rather than a pre-experiment estimate of the population size required for statistical significance.

### Visual stimuli

Full field illumination (diameter: 500-560 μm) was delivered to the preparation through a customized condenser from three LEDs (peak power at 405, 520 and 640 nm). Light intensities (photons/μm^2^/s) were converted to photoisomerization rates (R*/photoreceptor/s) using the estimated collecting area of rods and cones (1 and 0.37 μm^2^, respectively for primate; 0.5 μm^2^ for mouse rods), the LED emission spectra and the photoreceptor absorption spectra (Baylor et al., 1984; Baylor et al., 1987). In Figures 5 and 6 the short and long wavelength LEDs produced a mean of ~10-20 R*/rod/s and ~200 R*/L-cone/s. Rod-and cone-preferring flashes were 10 ms in duration.

Rods were selectively stimulated using the 405 nm LED. To test for cone contribution to these responses, we monitored the ratio of On parasol responses elicited by the 520 and 405 nm LEDs. This ratio is predicted to differ by a factor of ~2.4 for responses elicited by rods and M cones based on the rod and M cone spectral sensitivities (On parasol cells do not get significant S cone input (Field et al., 2010)). The ratio increased noticeably at 100-300 R*/rod/s, and increased by a factor of 1.8 at 300 R*/rod/s. Together with the theoretical factor of 2.4, this indicates that responses to the 520 nm LED represented a ~60% contribution from cones and ~40% from rods at 300 R*/rod/s. The ratio of rod sensitivity to cone sensitivity to the 405 nm LED is predicted to be 2.4-fold higher than that for the 520 LED, indicating that ~25% of the response to the 405 nm LED at 300R*/rod/s originated in cones. Cone contributions were substantially smaller at lower background light levels.

### Analysis

For spike recordings from ganglion cells, we detected spike times and compiled them into peristimulus time histograms as previously described (Murphy and Rieke, 2006). Assuming linearity, rod responses to sinusoidal stimuli (Figure 7) were predicted by convolving the measured flash responses with sinusoids of the various frequencies. Electrophysiology example traces presented throughout the figures represent the average of 5-20 raw responses to the same stimuli. All data are presented as mean ± SEM and two-tailed paired Student’s t-tests were used to test significance.

## Acknowledgements

We thank Cole Graydon for commenting on the manuscript, Chris English for assistance with tissue acquisition and Shellee Cunningham for general technical assistance. Support was provided by HHMI and NIH (EY028111).

---

**Figure 3–figure supplement 1:**
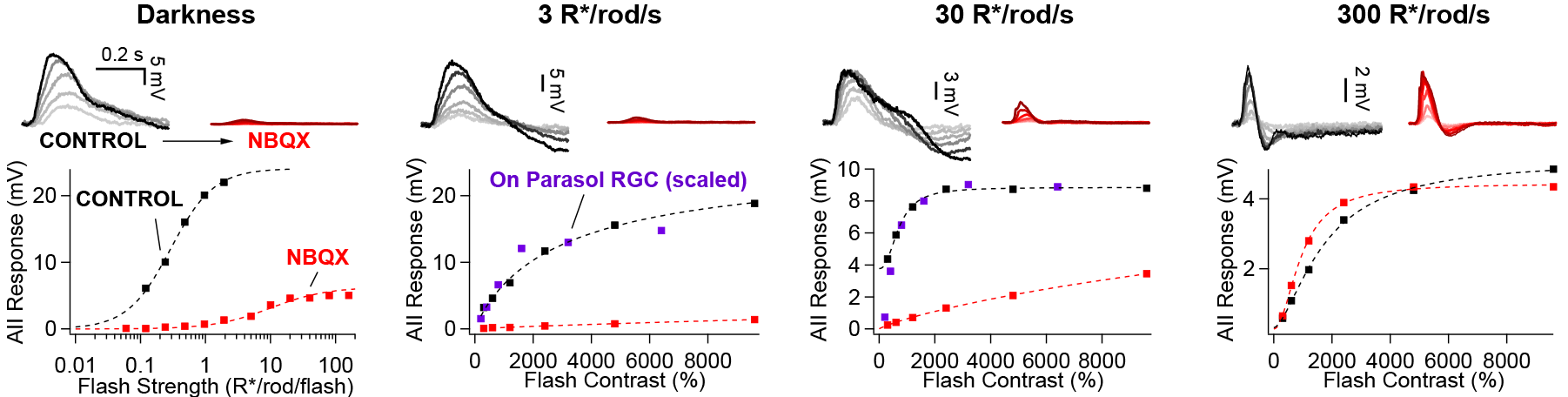
Rod signals in AII amacrine cells are blocked by NBQX (10 *μ*M) at low light levels. Example current-clamp recording of scotopic and mesopic signaling in an AII amacrine cell. Flash responses in darkness were largely eliminated by bath application of NBQX. NBQX-insensitive responses emerged as flash strength was further increased. In the presence of dim (3 R*/rod/s) and moderate (30 R*/rod/s) backgrounds, stimuli that produce sub-saturating responses in the RGCs (scaled) are almost entirely blocked by NBQX in AII amacrines. At a background of 300 R*/rod/s (i.e. above rod saturation) flash responses in AIIs are largely insensitive to NBQX application.

**Figure 3–figure supplement 2:**
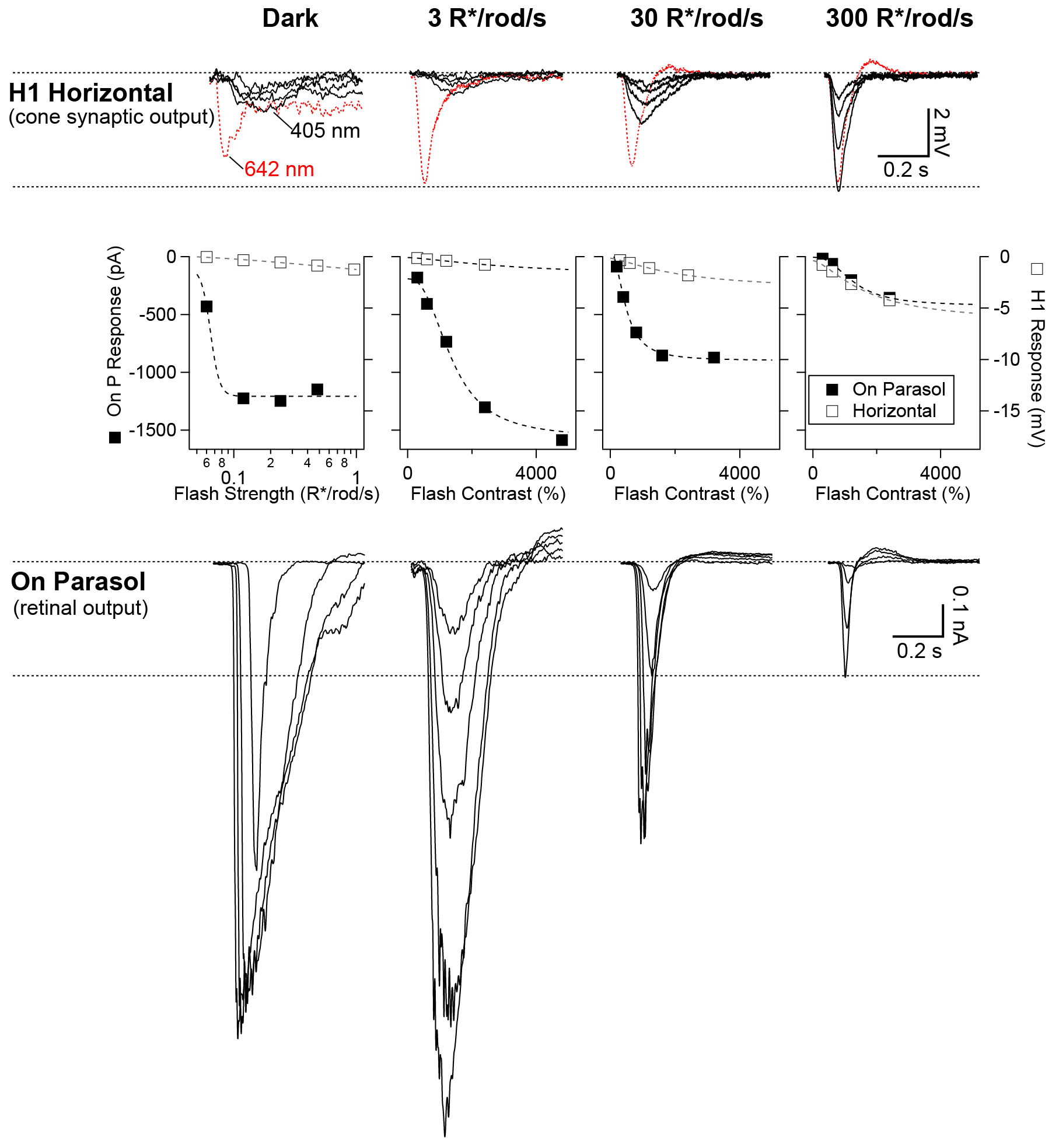
Rod signals are weak in H1 horizontal cells. Example recordings of an H1 horizontal cell (top row) and On parasol RGC (bottom row) from the same retinal mount. As in Figure 3–figure supplement 1, short wavelength rod-preferring (and occasional long wavelength cone-preferring) flash responses were recorded across a range of scotopic and mesopic backgrounds. Rod signals in H1 horizontal cells were weak at low and intermediate backgrounds, despite the cells” robust responses to long wavelength stimuli. At 300 R*/rod/s, long and short wavelength stimuli evoked responses with similar kinetics, suggesting they both arise from cones. On parasol RGC responses, however, were largest at the dimmest light levels. At 300 R*/rod/s, responses in On parasols had kinetics that were similar to those observed in horizontal cells. Right and left ordinate axes on the middle plots are scaled such the RGC and H1 responses are matched at 300 R*/rod/s (a background at which signals are dominated by cone activity).

**Figure 3–figure supplement 3:**
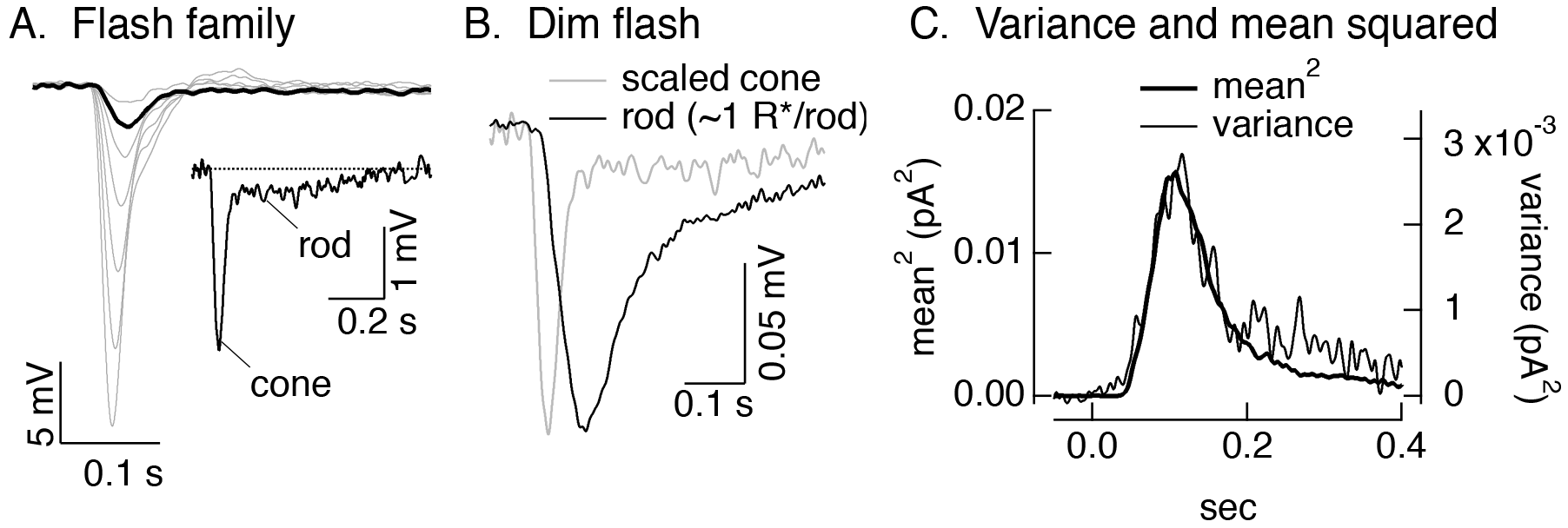
Rod responses measured in cones. A) Responses of a L cone to a family of brief 405 nm flashes. Inset shows the long “tail” associated with the rod response (see (Schneeweis & Schnapf, 1995a)). B) Average response of the same cell as in A to a flash producing ~1 R*/rod (black). For comparison, the gray trace shows the scaled response from the inset in A. C) Mean response squared and time-dependent variance (corrected for variance without a flash) for the same cell and responses in A and B. Assuming the variance is dominated by Poisson fluctuations in photon absorption, the ratio between the mean response squared and the variance provides an estimate of the mean number of absorbed photons contributing to the response. In this cell, that corresponds to a single photon response of 0.022 mV. For 6 cells, the mean was 0.046 ± 0.007 (mean ± sem), consistent with previous work (Hornstein et al., 2005).

**Figure 6–figure supplement 1.**
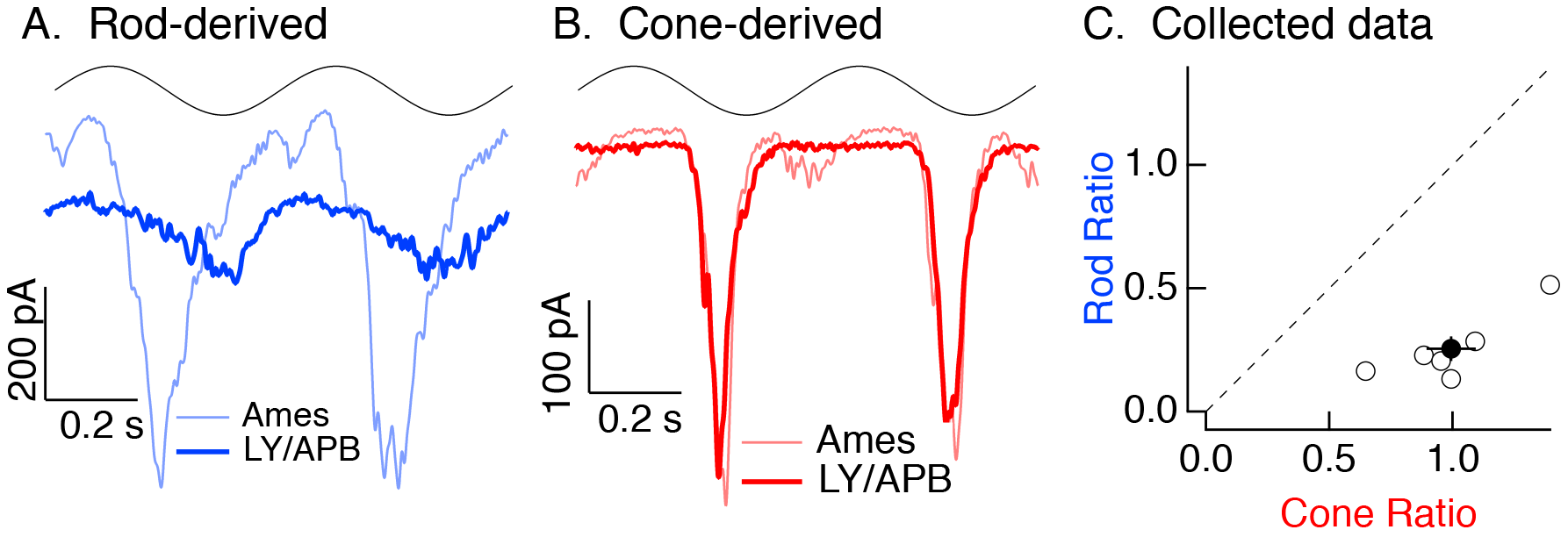
Rod, but not cone, signals in excitatory inputs to Off parasol RGCs arise from the primary rod pathway under mesopic conditions. Example recording of excitatory synaptic input to an Off parasol RGC in response to 2 Hz sine wave modulation of a short (rod-preferring) or long (cone-preferring) wavelength LED. Responses to short, but not long, wavelength light modulations were largely eliminated by a mixture of LY341495 and APB (see Methods).

